# Genomic evidence for rapid speciation and gene flow between sympatric carnivorous *Nepenthes* pitcher plants

**DOI:** 10.1101/2023.01.07.522986

**Authors:** M Scharmann, F Metali, T U Grafe, A Widmer

## Abstract

Speciation can result from both neutral and adaptive processes, but their relative importance and the factors exerting selective pressures are incompletely understood. In theory, interspecific gene flow could suffice to reverse speciation, or else erode neutral divergence and expose traits and underlying genes whose divergence is due to selection. Hence, introgression can shed light on selection during the speciation process. Here we study mixed assemblages of carnivorous *Nepenthes* pitcher plants, which frequently produce natural hybrids yet maintain distinct phenotypes. Using ddRAD-seq markers, we characterize divergence and introgression for eight *Nepenthes* species that grow sympatrically in communities of three to seven species at four locations in Southeast Asia, totalling 22 populations. The sympatric species fell into two discrete classes of high and low divergence. Five lineages with high divergence displayed little recent introgression in tests of location-dependent allele sharing (ABBA-BABA) despite the presence of some natural hybrids. However, all five lineages appear to have introgressed in the more distant past, as revealed by coalescent models with Approximate Bayesian Computation. In the same locations occur three further sympatric species with low genetic divergence. These incipient species also showed some natural hybrids, but in addition both ABBA-BABA tests and ABC suggested very recent or ongoing introgression, raising the question how divergence is maintained in these hybrid zones. One trait possibly involved in maintenance of divergence against gene flow might be the carnivorous pitcher traps, whose morphology showed greater divergence than expected under neutral evolution (Pst–Fst) in the introgressing species pair *N. hemsleyana* and *N. rafflesiana* t.f..

## Introduction

The degree of reproductive isolation (RI) and genetic divergence between populations and species can vary extensively, which is referred to as the speciation continuum (Seehausen et al. 2014). To disentangle the contributions of adaptive and neutral processes to the evolution of RI, speciation, and their dynamics (Nosil et al. 2017), different stages along this continuum need to be studied. While RI is complete at one end of the continuum, it is incomplete at the other end, where incipient speciation can be challenged by gene flow. Hybridization and repeated backcrossing between incipient species that have already acquired some divergence, for example during an earlier phase of isolation, can lead to introgression (Harrison and Larson 2014). Such gene flow can erode divergence over time, unless it is balanced by selection (Lenormand 2002). Without selection against at least some admixed genotypes (Lindtke et al. 2014; Christe et al. 2016), incipient species exposed to gene flow cannot sympatrically co-exist for long and the speciation process may be reversed (Vonlanthen et al. 2012). The stage along the speciation continuum where incipient species are maintained in the face of gene flow is also known as “isolation with migration” or “divergence with gene flow”, and encompasses hybrid zones formed by secondary contact or primary divergence (DGF, Pinho & Hey, 2010). Introgression, however, may have a dual role: in addition to challenging divergence between incipient species, it can also supply adaptive variation (Racimo et al. 2015; Suarez-Gonzalez et al. 2018), and even promote divergence and speciation (e.g. homoploid hybrid speciation: Gross & Rieseberg, 2005; hybrid swarm theory: Meier et al., 2017; Seehausen, 2004)(Seehausen 2004; Meier et al. 2017). These ‘modern’ views on speciation involving phases of incomplete RI and gene flow (Wu 2001) are in marked contrast to the ‘classic’ view (Mayr 1942) in which speciation proceeds only in the absence of gene flow, typically mediated by geographic isolation.

Hybrid zones provide the opportunity to empirically study the role of selection in ecological divergence and speciation, because the incipient species that persist in the face of gene flow are not able to accumulate or retain strong divergence unrelated to RI (Schluter 2000). Outlier scans for interspecific differentiation are widely used in such systems with the goal to identify traits and genomic regions underlying RI and speciation, and it is frequently assumed that outliers result from resistance to homogenizing gene flow. Through differential selection against migrant alleles, DGF can contribute to the formation of differentiation outlier loci and genomic islands of differentiation (Turner et al. 2005; Harrison 2012). However, such outliers can also arise from processes unrelated to RI and speciation *per se*, including divergence after speciation due to independent adaptation in descendant populations, background selection, and genomic variation in recombination rate (Cruickshank and Hahn 2014). To reject these alternative explanations in favor of diversifying selection and RI, it is necessary to verify the occurrence of interspecific gene flow.

Gene flow is difficult to uncover in practice. While early-generation hybrids are relatively easy to identify with molecular methods (Pritchard et al. 2000; Anderson and Thompson 2002; Vähä and Primmer 2006), they do not imply realized gene flow across species boundaries because these could fail to reproduce. Evolutionarily relevant gene flow, i.e. longer-term addition of foreign variants to a gene pool, must be inferred from current patterns of genetic variation. Instead of individuals of hybrid pedigree, samples representative of the gene pools are required. However, similar patterns of genetic variation can result from distinct evolutionary processes. Two conspicuous indicators of introgression are interspecific allele sharing and discordance of phylogenetic gene trees. However, interspecific allele sharing is likely to stem from retained ancestral polymorphism, because speciation events frequently occur much more rapidly than genetic coalescence occurs within natural populations. Gene tree discordance, on the other hand, can result from random sorting of such ancestral polymorphisms if they pass through several speciation events (Degnan and Rosenberg 2009). These ubiquitous confounding processes must be considered in inferences of gene flow between species.

A non-parametric class of introgression tests leverages the expectation that under a model of sequential population splits (or speciation events), any two sister lineages (and their further descendants) should be equally genetically different from lineages that split off earlier (e.g. (Durand et al. 2011; Pease and Hahn 2015; Blischak et al.). This expectation also implies that the genetic divergence between two species is always the same regardless of which sub-populations of the species are compared. Along this line of reasoning, reduced Fst between sympatric populations of two species, as compared to Fst between allopatric populations, was interpreted as a signature of introgression in several examples, e.g. *Heliconius* butterflies (Martin et al. 2013), or subspecies of *Zea mays* (Hufford et al. 2013). A caveat of the Fst metric is that it is sensitive to demography. Thus, historical differences in effective population size (i.e. genetic drift), which are likely to occur during range expansions, could suffice to explain the sympatric-allopatric Fst differences. A statistic that essentially exploits the same expectation as the sympatric-allopatric Fst comparison, but is robust to demographic differences, is the D- or ABBA-BABA statistic (Durand et al. 2011). This statistic measures allele sharing that is not expected under a model of pure drift (and mutation), and was originally developed to test for introgression between non-African humans and Neanderthals. However, the statistic may also be configured to test whether interspecifc allele sharing in sympatry is higher than in allopatry, considering drift and mutation along an unrooted phylogenetic tree of two species with two-sub-populations each. Although these methods are effective at discerning introgression from confounding processes (see above), they typically provide limited insights on the timing, direction, magnitude and other detailed aspects of introgression.

Another class of methods to infer introgression employs parameterized evolutionary models (e.g. (Hey and Nielsen 2007; Roux et al. 2013; Lohse and Frantz 2014; Dalquen et al. 2017), which can allow much more detailed inferences. In general, these models consider speciation as a population process and predict distributions of gene trees or population genetic summary statistics. Heuristics can then be used to fit the models to observed data, and parameters of introgression can be inferred. A frequently used class of strategies is the combination of coalescent theory (Rosenberg and Nordborg 2002; Wakeley 2009) with simulations and Approximate Bayesian Computation (Tavare et al. 1997; Beaumont et al. 2002). This framework is particularly powerful for detecting introgression and estimating parameters, and recently critically important principles were implemented (Roux et al. 2013; Roux, Fraisse, et al. 2014; Roux et al. 2016) that were lacking in previous, more error-prone applications (Cruickshank and Hahn 2014).

Many studies of introgression investigated species pairs with geographically defined hybrid zones (Barton and Hewitt 1985; Anderson and Thompson 2002; Buerkle 2005; Gompert and Buerkle 2011; Abbott 2017). The hybrid zone framework offers great conceptual advantages to detect introgression and diversifying selection, but it is not straightforward to apply when more than two species are involved and when ‘pure’ parental reference populations are not spatially manifest. Genomic studies testing introgression among more than two sympatric species are rare. Only 12 out of 42 studies reviewed by Payseur & Rieseberg (2016) investigated more than two at least partially sympatric species. Recent studies of groups that might involve ongoing gene flow between multiple species include lake-specific cichlid radiations (Meyer et al. 2016; Meier et al. 2017), North American *Quercus* (Eaton et al. 2015) and *Populus* (Chhatre et al. 2018), *Pedicularis* (Eaton and Ree 2013), *Heliconius* butterflies (The *Heliconius* Genome Consortium 2012; Martin et al. 2013), *Anopheles* spp. (Fontaine et al. 2015), and Darwin’s finches (Lamichhaney et al. 2015). One explanation for this relative shortage of examples may be a lack of appropriate methods to study introgression in sympatric species groups, although these may be common in nature.

Here we test for introgression and divergence in the face of gene flow in sympatric assemblages of closely related species, with replicated sympatry at multiple geographic locations. We configure the D-statistic (Durand et al. 2011) to test for excessive interspecific allele sharing in sympatry, and develop an ABC approach to distinguish ancient introgression from introgression that is ongoing in the extant plant communities.

Our study system are sympatric communities of carnivorous *Nepenthes* pitcher plants, in which frequent observations of fertile hybrids indicated incomplete RI and raised the question how species may be maintained in the face of gene flow. *Nepenthes* (Nepenthaceae, Caryophyllales) contains c. 160 species of perennial carnivorous vines and shrubs (Cheek and Jebb 2001; Clarke et al. 2018). Botanical carnivory in general is understood as a nutrient sequestration strategy, supplying minerals lacking from soils (Ellison 2006). *Nepenthes* plants capture, drown and digest animals in their pitcher traps, which are highly modified leaves with complex biomechanical and physiological adaptations (Juniper et al. 1989; Moran and Clarke 2010; Ellison and Adamec 2017). Within the genus, there is limited variation in flower, inflorescence architecture and vegetative traits, but tremendous variation in pitcher trap size, shape, coloration, life span, and multiple other characters. However, no satisfactory explanation has yet been put forth for this diversity in what was proposed to represent an adaptive radiation (Meimberg and Heubl 2006; Bauer et al. 2012; Pavlovič 2012). One hypothesis is that carnivory itself promoted diversification through competition for prey, pushing coexisting species to exploit different prey spectra (Moran et al. 1999; Bauer et al. 2011; Chin et al. 2014; Gaume et al. 2016). This hypothesis implies that diversifying selection acts on trap morphology.

Here we aim to elucidate speciation processes in *Nepenthes* by studying signatures of past and ongoing introgression in sympatry. Under long-term introgression, neutral genes and traits are expected to move freely between species, while genes and traits under diversifying selection may continue to diverge. In other words, we hypothesize that interspecific gene flow among sympatric *Nepenthes* could indirectly reveal a role for divergent selection. First generation natural hybrids among sympatric *Nepenthes* are well documented at the phenotypic level (Cheek and Jebb 2001; Clarke 2001; Peng and Clarke 2015) and motivated this study. Hybrids demonstrate that RI is incomplete, but it currently remains unknown whether they mediate realized gene flow between species.

We focused on the inference of gene flow in the particularly rich *Nepenthes* assemblages in the lowland peat swamp and heath forests of Borneo, where seven or more species may grow within a few hundred meters distance (Figure 1). Three species were also studied in Singapore. No prior study had surveyed nuclear genetic variation in these *Nepenthes*. Specifically, we addressed the following questions: 1) Are sympatric *Nepenthes* morphospecies genetically distinct, and how are they related to each other? 2) What is the evidence for past and ongoing introgression between them, and at what stages of the speciation continuum are they? 3) Is divergence in *Nepenthes* pitcher trap morphology higher than expected under neutral evolution?

**Figure 1.**
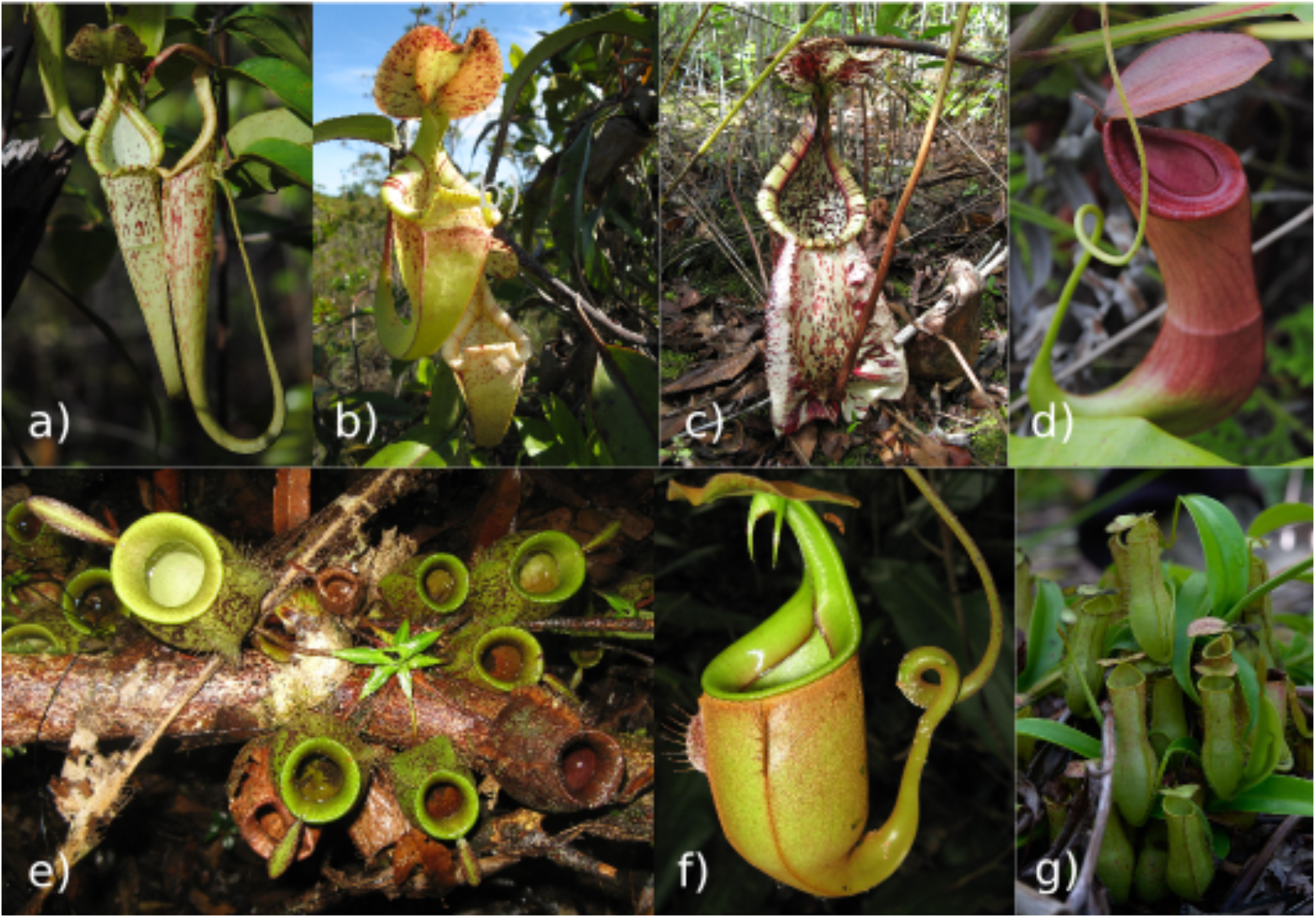
Portraits of sympatric *Nepenthes* morphospecies investigated in this study. a) *N. hemsleyana*, upper pitchers; b) *N. rafflesiana* typical form (t.f., Borneo endemic), upper pitchers; c) *N. rafflesiana* giant form (g.f., Borneo endemic), lower pitcher; d) *N. mirabilis*, upper pitcher (“typical” variety) e) *N. ampullaria*, lower pitchers; f) *N. bicalcarata*, upper pitcher; g) *N. gracilis*, lower pitchers. Photos: M. Scharmann.

## Results

### population genetic structure and natural hybrids

In our sample of 22 *Nepenthes* populations from four locations in Southeast Asia (Brunei, Binyo and Kuching on the island of Borneo, and Singapore), we found overall strong agreement between morphospecies and genetic clusters, using NJ trees (Figure 2), PCAs (Figures S3), and fastSTRUCTURE (Figures S4). Fifteen (5%) phenotypically inconspicuous individuals fell in between the main clusters in NJ trees and PCAs and were subsequently identified as F1, F2 and backcross hybrids (Table S5), while all other individuals (280) were assigned to a single genetic cluster with >99% probability (fastSTRUCTURE, Figures S4). Most populations contained few hybrids, except *N. rafflesiana* g.f. from Brunei where half of all individuals were identified as hybrids. All hybrids were excluded from further analyses, because we were interested in long-term introgression and not in hybridization events, which may be linked to recent deforestation.

**Figure 2.**
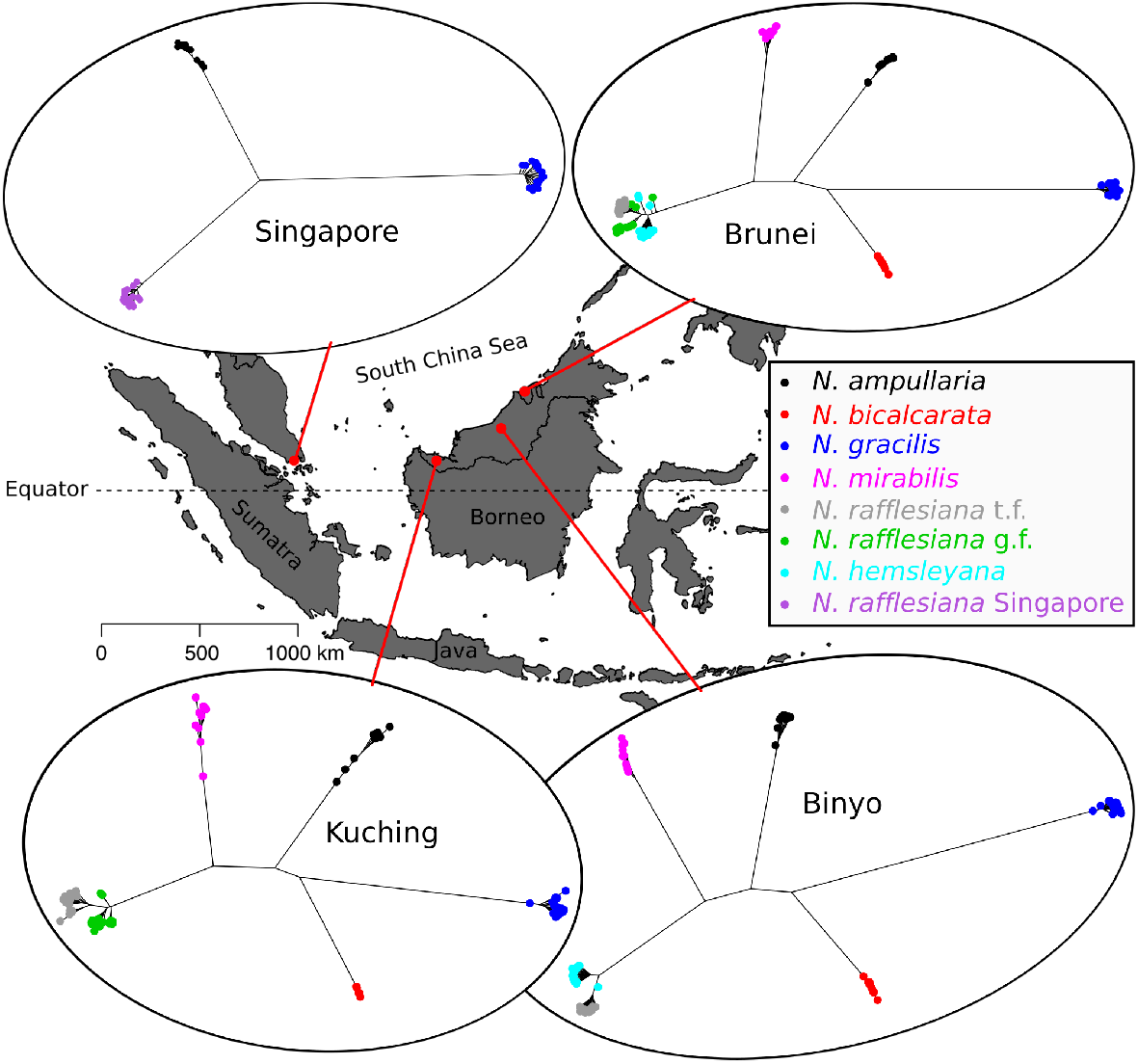
Sympatric *Nepenthes* sampled at four locations in Southeast Asia and clustered by genetic distance (NJ trees). 295 individuals were sampled in three countries (Brunei, Malaysia and Singapore). Morphospecies as identified in the field are color-coded, and genetically identified natural hybrids (Table S5) are included here with their erroneous field identifications.

### Population phylogeny, and patterns of genetic divergence and diversity

Considering all 22 *Nepenthes* populations together across the four geographic locations in a phylogenetic tree, the morphospecies grouped together as monophyletic clades, rather than by geographic location (Figure 3, top). However, the explanatory value of this phylogenetic model using a concatenated supermatrix alignment may be limited, because it ignores variation in geneaologies and polymorphism within lineages. Indeed such assumptions were violated in our data, as about half of all polymorphisms were shared between at least two species, and while the other half was species-specific, only a very small fraction (∼ 2%) of polymorphisms were species-specific and fixed in any species (Table 1).

**Table 1.** Fixed and shared polymorphisms between *Nepenthes* species after hybrid exclusion. Shown are percentages of 12,366 total SNPs, in 1,788 RAD-tags. The “shared SNPs” of a species are global SNPs whose alleles are found in at least one further species.

| species | N individuals | N populations | species-specific SNPs | fixed species-specific SNPs | shared SNPs |
| --- | --- | --- | --- | --- | --- |
| *N. gracilis* | 76 | 4 | 17.5 | 1.2 | 82.5 |
| *N. bicalcarata* | 22 | 3 | 5.3 | 0.2 | 94.7 |
| *N. ampullaria* | 33 | 4 | 13.8 | 0.5 | 86.2 |
| *N. mirabilis* | 27 | 3 | 10.6 | 0.3 | 89.4 |
| *N. rafflesiana* Singapore | 17 | 1 | 1.8 | 0 | 98.2 |
| *N. rafflesiana* giant form | 21 | 2 | 0.3 | 0 | 99.7 |
| *N. hemsleyana* | 33 | 2 | 1.2 | 0 | 98.8 |
| *N. rafflesiana* typical form | 52 | 3 | 1.1 | 0 | 98.9 |
| **overall** | **281** | **22** | **51.6** | **2.3** | **48.4** |

**Figure 3.**
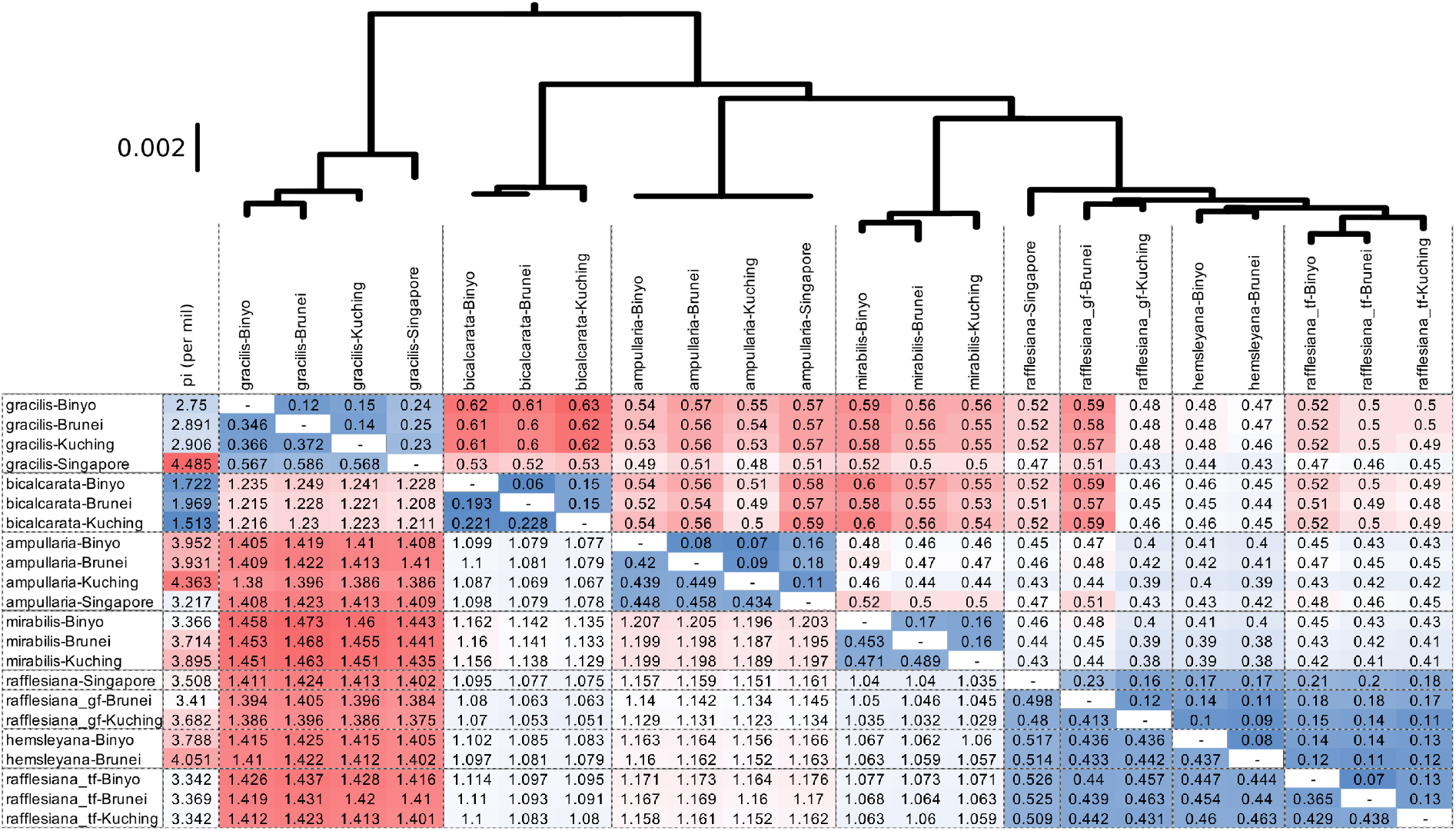
Maximum-likelihood phylogenetic tree (top), and selected diversity and divergence statistics for 22 *Nepenthes* populations and their pairwise combinations (below). First column: nucleotide diversity pi (per mil), lower triangle: absolute divergence dxy (average pairwise nucleotide differences, percent), upper triangle: Fst (Weir & Cockerham 1984). Color scales (from blue, lowest, over white to red, highest) are calculated within each of pi, Fst and dxy. Natural hybrids (Table S5) were excluded. Tree and table are derived from the same data (12,366 SNPs and 165,313 invariant sites).

At the level of populations, the *Nepenthes* showed threefold variation in nucleotide diversity (Figure 3, first column). *N. bicalcarata* populations were consistently the least diverse, and *N. gracilis* in Singapore was the most diverse population. Overall, the genome-wide average divergence between populations of the same species ranged from Fst 0.06 to 0.25, and dxy 0.193% to 0.586%. The low and high extremes of intraspecific divergence were again marked by *N. bicalcarata* and *N. gracilis*, respectively. Interspecific divergence ranged from Fst 0.1 to 0.63 and dxy 0.44% to 1.47% (Figure 3, bottom), notably overlapping with levels of intraspecific divergence.

Among the species, two classes of interspecific divergence levels were eminent (Figure 3, bottom): a high-divergence class consisting of the four species *N. gracilis, N. bicalcarata, N. ampullaria* and *N. mirabilis*, and a low-divergence class encompassing *N. rafflesiana* Singapore, *N. rafflesiana* t.f., *N. rafflesiana* g.f. and *N. hemsleyana*. The latter four species form a complex of closely related species, which we subsequently refer to as the *N. rafflesiana* species complex. The high-divergence lineages generally showed dxy >1% and Fst c. 0.4 or greater, whereas low-divergence species of the *N. rafflesiana* complex showed dxy <0.6% and Fst <0.23. The two classes of divergence are also reflected in the branch lengths of NJ trees (Figure 2), the PCA coordinates (Figures S3), the branch lengths of the population phylogeny (Figure 3, top), and the number of fixed, species-specific SNPs (Table 1).

### Introgression in sympatric species pairs tested with Patterson’s D-statistic

We hypothesised that species pairs may have more alleles in common at one location than at another. Such excessive allele sharing may indicate introgression, which we evaluated with the D-test, leveraging our geographic sampling scheme with species pairs occurring together at two (or more) locations (see Introduction, Figure 2). Under our sampling, 43 D-tests were possible. Nine tests revealed significant D-statistics (Table S6), indicating that introgression had occurred in sympatry between different morphospecies. Introgression in sympatry was found for eight of the 21 species pairs in lowland Borneo and Singapore and encompassed all studied species. The strongest D-statistics were found between members of the *N. rafflesiana* complex, namely for the pairs *N. rafflesiana* g.f. - *N. rafflesiana* t.f., and *N. hemsleyana* - *N. rafflesiana* t.f. The D-statistic cannot distinguish at which of the geographic locations introgression occurred, but the introgressed populations are expected to show lower genetic divergence. Accordingly, introgression between *N. rafflesiana* g.f. - *N. rafflesiana* t.f. probably occurred in Kuching (dxy Kuching 0.431%, dxy Brunei 0.439%, Figure 3 bottom), and between *N. hemsleyana* - *N. rafflesiana* t.f. in Brunei (dxy Brunei 0.440%, dxy Binyo 0.447%, Figure 3 bottom).

### Introgression tested by coalescent models and ABC

Complementary to the D-tests and to gain further insights into introgression, we used a coalescent framework that considered confounding processes such as retention of ancestral polymorphisms, changes in population sizes (demography), background selection, and genomic as well as temporal variation in introgression rates. For each of 54 sympatric species pairs, we performed extensive simulations and ABC model choice between four models that differed in the level and timing of introgression (Figure 4).

**Figure 4.**
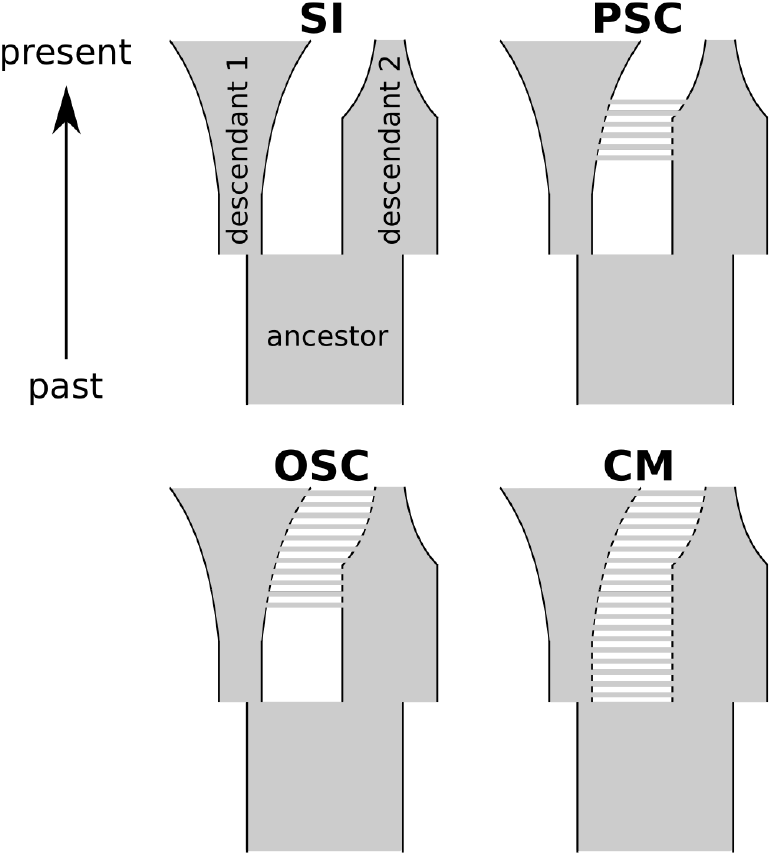
Four coalescent models of speciation were explored for sympatric *Nepenthes* species pairs. Clockwise from top left: strict isolation (SI), past secondary contact (PSC), continuous migration (CM), and ongoing secondary contact (OSC). In SI, an ancestral species instantly splits into two completely isolated descendants. CM follows the same course, but the descendant species exchange migrants at a constant rate since their split and continuously up to the present. In PSC, species are first isolated but then exchange genes during a limited phase of secondary contact, which ceases before the present. OSC differs from PSC by the extension of migration up to the present. In these example schemes, that show just one parameter combination of independent population sizes and changes under each of the four models, descendant species 1 had a small initial population size just after speciation but then underwent exponential growth during a long phase up to the present, while descendant species 2 was initially demographically larger but recently underwent exponential population decline.

In a first step, we asked whether there was any introgression at all, by competing speciation histories without introgression (model SI) against those with introgression (aggregated models PSC, OSC and CM). ABC model choice with ABC-RF (Pudlo et al. 2016) showed high power for this classification (prior error rates 10.5 - 13.6%, Table S7). Introgression was significantly more probable than strict isolation (SI) for 51 out of 54 sympatric *Nepenthes* species pairs (Bayes Factor > 3, Figure 5, filled black circles), and only three sympatric pairs remained ambiguous about introgression (Figure 5, blank circles).

**Figure 5.**
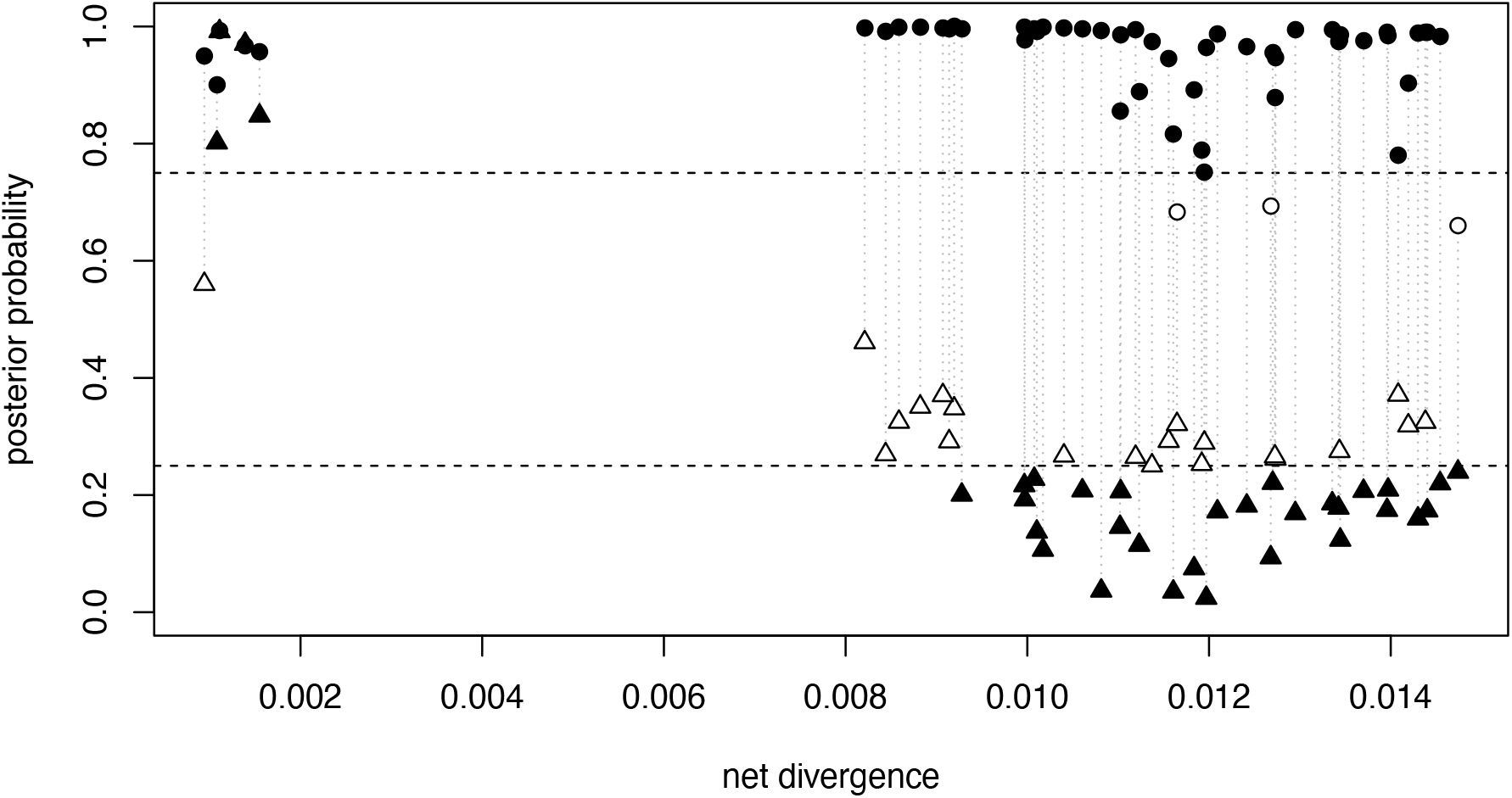
Posterior probability (PP) of introgression in 54 sympatric *Nepenthes* species pairs, plotted in relation to their net sequence divergence (per-site interspecific divergence dxy minus intraspecific diversity pi, (Nei and Li 1979)). Each species pair was classified twice with ABC model choice, and dotted vertical lines connect the two results for each species pair. Circles indicate the PP for any introgression at all (ABC-RF model choice of aggregate models PSC, OSC and CM versus SI). Triangles indicate the PP for the timing of introgression, i.e. high PPs suggest that introgression is ongoing while low PPs indicate that it occurred in the past but has since ceased (ABC-RF model choice of aggregate models OSC and CM versus PSC). Filled symbols indicate results with Bayes Factor >= 3, while blank symbols show results in the “ambiguity zone” marked by horziontal lines at PP 0.25 and 0.75. Details see Table S7.

These all involved populations of *N. bicalcarata* (Table S7; *N. bicalcarata* against *N. mirabilis* at Binyo, against *N. ampullaria* at Kuching, and against *N. gracilis* at Kuching). However, introgression was significantly more probable than SI for the same species pairs at other locations. Thus, all investigated *Nepenthes* species revealed signatures of introgression and this encompassed all 21 species combinations.

Notably, there was no simple relationship between interspecific divergence (net divergence) and support for introgression (Figure 5, circles). Introgression was highly probable (PP >= 0.9) for all species pairs with net divergence lower than 1.1%, while the 33 species pairs with net divergence >= 1.1% in eleven cases recieved lower, yet in all but three cases still conclusive, probabilities for introgression (see above, Figure 5, Table S7). All of these lower-support cases involved N. *gracilis* and/or N. *bicalcarata*, the two most strongly diverged species. However, most of the cases with lower probabilities for introgression (ten out of eleven) did not involve the most diverged species N. *gracilis* (avg. net divergence over 18 species pairs 1.37%, Table S7, Figure 3), but rather the less diverged N. *bicalcarata* (avg. 0.6 net divergence over 16 species pairs 1.23%, Table S7, Figure 3). Thus, support for introgression was not generally lower for species with higher divergence.In a second step, we asked when in time introgression had occurred. Specifically, we performed ABC-RF model choice to test whether gene flow was still ongoing up to the present (aggregated models CM, OSC), or alternatively, had stopped at some point in the past (PSC; Figure 5, triangles). Statistical power for this distinction was good with prior error rates of 10.3 - 14.1% (Table S7). For 29 of 54 species pairs, ongoing gene flow (CM, OSC) was rejected in favor of a temporally limited, past phase of introgression (PSC). These species pairs with old introgression were all in the high-divergence class at net divergence >0.9%. In species pairs with lower divergence, ongoing gene flow was slightly more probable, mainly for *N. mirabilis* against the *N. rafflesiana* complex (net divergence 0.8 - 0.95%, Table S7), yet these results were ambiguous (Bayes Factors < 3, Figure 5, empty symbols). Overall, there were 21 sympatric pairs spanning the entire range of divergence levels, in which models could not distinguish between ongoing and past introgression. Clear support for ongoing introgression was obtained only in the low-divergence class of species (net divergence < 0.16%), involving four of the five species pairs in the *N. rafflesiana* complex. At Brunei, where three members of this group are sympatric, ongoing gene flow was supported for *N. rafflesiana* t.f. with both *N. hemsleyana* and *N. rafflesiana* g.f. (Table S7). At Binyo, ongoing gene flow was detected for *N. rafflesiana* t.f. - *N. hemsleyana*, and at Kuching for *N. rafflesiana* t.f. - *N. rafflesiana* g.f. (Table S7).

For these four introgressing species pairs we attempted to further disentangle the history of introgression, asking whether they represent secondary contacts (OSC) or cases of primary divergence with gene flow (CM). However, the two models were in general poorly distinguishable by ABC-RF (prior error rates ranged from 38.8 - 41.3%), and the posterior probabilities for any of the four species pairs remained ambiguous (Table S7). Thus, both secondary contact and primary divergence with gene flow remain plausible scenarios for the origin of the three Bornean species in the *N. rafflesiana* complex.

### Pst–Fst comparison for *N. hemsleyana* and *N. rafflesiana* t.f

For one population pair of *N. hemsleyana* and *N. rafflesiana* g.f. that grows in sympatry at one site in Brunei, hence in a “natural common garden”, we estimated pitcher trait divergence by the Pst metric and tested whether it exceeds neutral divergence, represented by the genome-wide distribution of Fst. We found that divergence of functionally important pitcher shape traits, including the relative distance between peristome and “girdle” (the waxy zone or tubular section) and the relative internal and external diameters of the pitcher orifice (Figure 6, Table S8), are strongly elevated over zero and incompatible with the null expecation under drift. Similar signatures of elevated phenotypic divergence were also found for pitcher wall thickness, the relative external orifice diagonal (perpendicular to the other orifice measurement), and absolute pitcher length. However, for the latter two traits, our approach lacked robustness, as indicated by c/h2 values larger than 0.2 (Brommer 2011).

**Figure 6.**
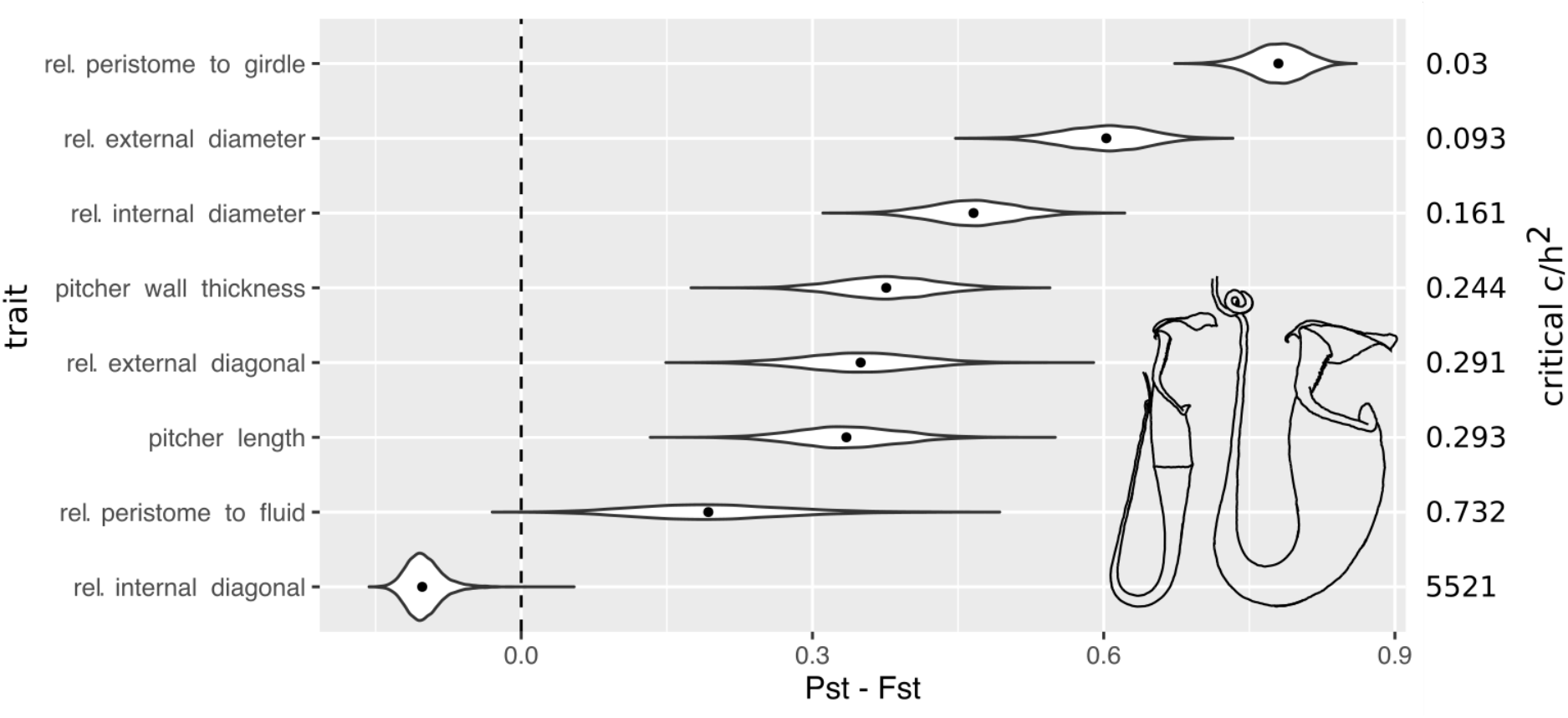
Comparison of observed phenotypic divergence (Pst) against neutral divergence (Fst, genome-wide mean = 0.11) in upper pitcher trap traits of the sister species *N. hemsleyana* and *N. rafflesiana* t.f.. Shown are the full distributions (violins, means as dots) of the Pst–Fst difference (X-axis) for 10,000 bootstrapped combinations of phenotypes and genetic markers under the assumption that c/h^2^ = 1, and the critical c/h^2^ above which the 95% confidence intervals of Pst and Fst do not overlap. The lower the critical c/h2, the more robust the assumption of c/h2 = 1 and thus the suitability of Pst to approximate Qst. Insets in the lower right corner illustrate the divergent upper pitcher shapes in lateral view, scaled to same length, of *N. hemsleyana* (left) and *N. rafflesiana* t.f. (right).

Pst-Fst distributions for the relative distance between peristome and digestive fluid level and for the relative internal orifice diagonal included zero, and were thus compatible with the null expectation (Figure 6).

## Discussion

The *Nepenthes* morphospecies studied here correspond to distinct genetic clusters in sympatry (Mallet 1995), and their populations sampled at different locations grouped by morphology in a phylogenetic tree. We thus show that these *Nepenthes* morphospecies, understood as testable hypotheses, represent distinct evolutionary lineages. A bifurcating species tree, however, falls short of explaining population genomic patterns in these species. One noticeable violation of tree-like evolution is that *N. gracilis*, as the most basal lineage, did not show uniform divergence (dxy) against all other species. Instead, divergence was reduced against *N. bicalcarata, N. rafflesiana* g.f., and *N. ampullaria* at Kuching (Figure 3, bottom). Another example is that *N. ampullaria* showed higher divergence against *N. mirabilis* than against *N. bicalcarata* and the *N. rafflesiana* complex. Under tree-like evolution, *N. ampullaria* should be most divgerged from *N. bicalcarata*, and lower and similar against *N. mirabilis* and the *N. rafflesiana* complex.

In comparison to other congeneric plant species for which similar genome-wide estimates of divergence are available, the *Nepenthes* range at low to intermediate levels (dxy c. 0.4 to 1.5%). As points of reference, the two sympatrically derived species of *Howea* palms show dxy 0.2% (Dunning et al. 2016), whereas the hybridising herbs *Geum urbanum* - *G. rivale* dxy 1.1% (Jordan et al. 2018), and the hybridising trees *Populus tremula* - *P. alba* have a dxy of 2.7% (Christe et al. 2017). It should be noted that RAD-like markers as used here may lead to somewhat biased estimates of diversity and divergence (Arnold et al. 2013; Gautier et al. 2013), and therefore our estimates are best understood as relative rather than absolute values. However, we minimised the effect of null alleles (allele drop-out) by considering only loci that were present at high frequency (present in 70% or 80% of individuals of the considered species), and we reduced the bias for polymorphic genome regions by the inclusion of invariant RAD-tags.

This study provides several lines of evidence that, despite their genetic distinctness, all of these Nepenthres species have histories of introgression (Figure 7), consistent with the long known presence of natural hybrids ((Cheek and Jebb 2001; Clarke 2001; Peng and Clarke 2015); Table S5). First, the observed variation in interspecific allele sharing by location (ABBA-BABA tests) indicates introgression which happened *after* the extant spatial distributions of *Nepenthes* were established (Figure 7, Table S6). To interpret the D-test results and the age of the introgression signatures which it assesses, it is crucial that the D-tests were based on differences between geographically defined populations of the same species. Thus, the age of current *Nepenthes* spatial distribution gives an upper boundary for the age of these introgression signatures. Under the assumption that *Nepenthes* distribution is associated with the tropical warm climate vegetation of coastal heath and peat swamp forests, the current distribution has established following the Last Glacial Maximum (∼21 ka BP: Hunt, Gilbertson, & Rushworth, 2012; Raes et al., 2014), and is likely younger than the marine regression that exposed some of our sampling locations approximatley ∼15 ka BP (Dommain et al. 2014). The peat domes from which our Brunei populations were sampled started to grow at only ∼2.8 ka BP (Dommain et al. 2015). Consequently, the introgression inferred from D-tests is likely to have occurred during the last few hundred, but maximum 2,000 generations, if a mean generation time of 10-15 years is assumed, and can be considered evolutionarily recent.

**Figure 7.**
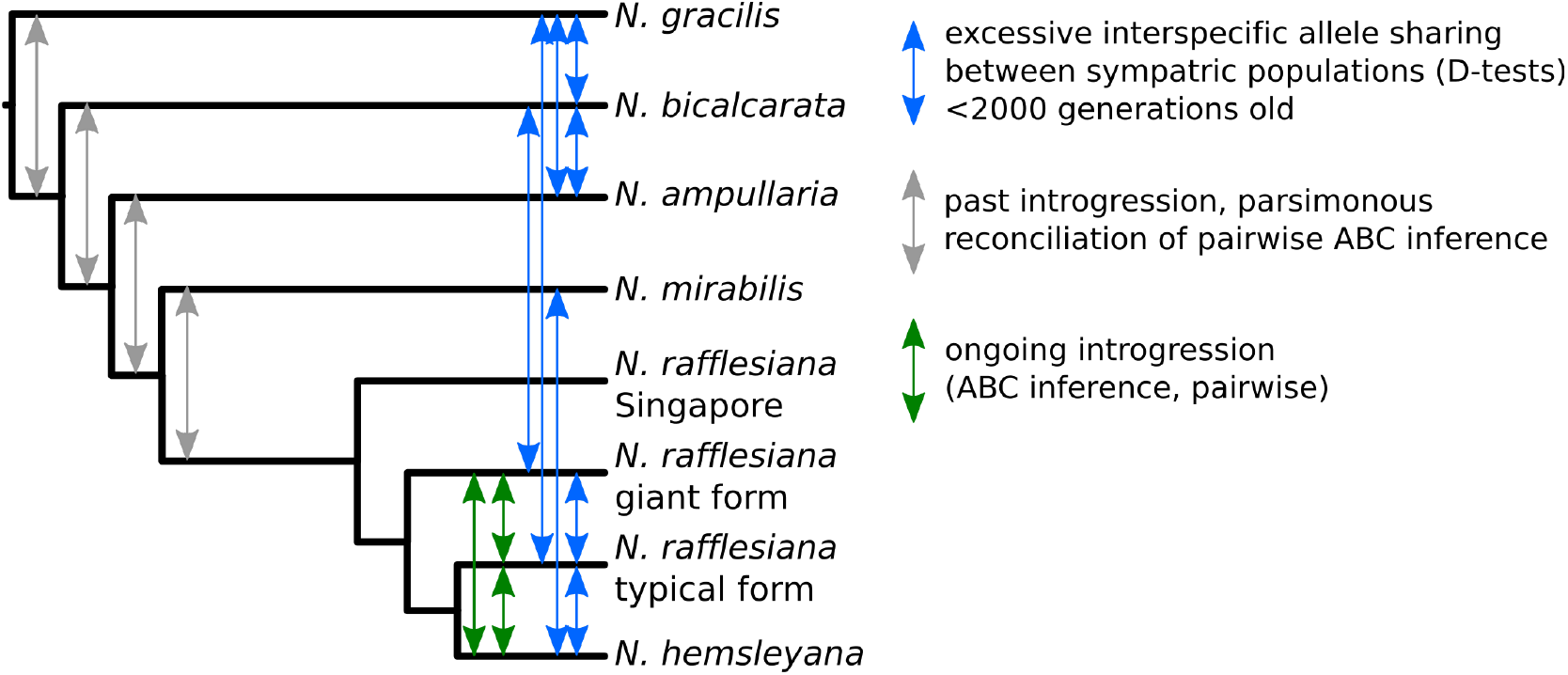
Bifurcating species tree hypothesis and schematic summary of introgression signatures detected between *Nepenthes* species. Branch lengths not to scale.

Our second strategy using coalescent modelling and ABC suggested introgression for all investigated *Nepenthes* lineages, and allowed to further characterize the timing of introgression as either ancient, or ongoing (Figure 7, Figure 5, Table S7). Methods for testing migration between demes, corresponding to introgression between species, have continuously been improved in recent years. For example, a simple scenario of long-term gene flow between species that maintain species boundaries can now be investigated across many independent loci, where effective gene flow is expected to be variable along the genome (Wu 2001; Roux, Fraïsse, et al. 2014; Aeschbacher et al. 2017), corresponding to a semi-permeable species boundary (Harrison and Larson 2014). The coalescent models we developed here incorporate such variation in migration rates between loci, and consider three relevant confounding factors together, which are (1) demographic disequilibrium (population expansion or shrinkage), (2) variation in duration and timing of migration, and (3) genomic variation in Ne due to background selection (Charlesworth et al. 1997; Ewing and Jensen 2016; Roux et al. 2016). Especially the modelling of temporal variation in migation rates allowed crucial inferences in this study. Furthermore, we explored several novel population genetic summary statistics, some of which proved highly informative, e.g. moments of genome-wide distributions of Fst, net divergence and dxy (5th percentile, mean content of the three lowest bins in a histogram), the interspecific correlation coefficient of nucleotide diversity, and joint site-frequency spectra, which are gaining popularity in ABC modelling of speciation (Xue and Hickerson 2015; Smith et al. 2017). An improved ABC model choice procedure that employs machine learning (Pudlo et al. 2016), which is robust to common statistical problems such as the “curse of dimensionality” and un-informative summary statistics (e.g. <u>Blum et al. 2013</u>), gave us the power to distinguish between absent, ancient, and ongoing introgression even for species pairs whith overall low divergence, and in the presence of complex confounding factors.

We note that the D-tests and ABC inference were not always consistent in regard to introgression. D-tests are non-parametric and based on the comparison of four populations, while our coalescent models considered pairs, which is an advantage because fewer populations are required, but apparently less sensitive than the D-tests to some signatures of introgression. The D-tests as configured here, however, can not detect introgression that is older than geographic population structure, and can not reveal whether introgression is ongoing. In particular, there were cases in which D-tests indicated recent introgression but ABC failed to support ongoing introgression. These may be cases in which introgression started less than 2,000 generations ago (explaining significant D-tests, see above), but then stopped before the present (explaining lack of support for ongoing introgression from ABC). Alternatively, ongoing introgression may have recieved a low posterior probability because signatures of recent introgression were weaker than signatures of past introgression in the forced bipolar ABC model choice - yet both are in reality not mutually exclusive.

The sympatric *Nepenthes* species pairs investigated in this study fall into two distinct classes of evolutionary divergence and associated gene flow histories (Figure 5, Figure 7), rather than a single class or a speciation continuum (Seehausen et al. 2014). Our low-divergence class comprises three members of the *N. rafflesiana* complex that grow sympatrically in Borneo, and F1 as well as backcross hybrids are found naturally. However, these contemporary hybrids were not responsible for signatures of gene flow, because clear signals were obtained after hybrid exclusion. This suggests that natural hybridization does proceed further than early, discernable hybrid generations and indeed leads to introgression. For this species complex, both D-tests and ABC inference supported recent and ongoing introgression at each of three geographic locations in Borneo, and possibly simultaneous among all three species where they co-occur. Notably, the ABC inference of ongoing gene flow is robust to confounding factors (see above) and independent of the history of isolation, i.e. it does not matter whether these hybrid zones are assumed to originate as secondary contacts or from primary divergence with gene flow. Indeed these two scenarios (formalised as the models OSC and CM, Figure 4) were indistinguishable by our methods. Despite evidence for introgression, the phylogenetic species tree grouped the morphospecies as distinct evolutionary lineages with single origins (Figure 3, Figure 7), suggesting that species boundaries are maintained in sympatry and divergence continues in the face of gene flow. The Singaporean member of the N. *rafflesiana* species complex is not sympatric with other species from the complex, and since geographic isolation alone may suffice to explain its divergence we do not discuss it further here.

Fully sympatric with members of the *N. rafflesiana* complex, which form the low-divergence class, are the four species *N. ampullaria, N. bicalcarata, N. gracilis*, and *N. mirabilis*. These species are more divergent from each other and the *N. rafflesiana* complex. Natural early generation hybrids have been reported for all possible combinations of these four species (Clarke, 1997; pers. obs. M. Scharmann) but are less common than between members of the *N. rafflesiana* complex. Consistently, only few of their populations revealed signs of recent introgression in the D-tests. The more elaborate ABC inference supported introgression between all five lineages, but this introgression was in most cases probably confined to the more distant past. The few ambiguous results of ABC model choice for past versus ongoing introgression could be due to introgression that was not ongoing anymore but that stopped not very long ago, thus displaying population genetic patterns intermediate between the expected outcomes of the past and ongoing introgression models.

Regarding our result that most introgression between high-divergence lineages was not recent, it is important to acknowledge that our pairwise tests (ABC) are not phylogenetically independent. We argue that past introgression detected with multiple descendants of a phylogenetic lineage is most parsimoniously explained as a single introgression phase between the ancestral lineages, rather than independent events between all descendants. When the 46 signatures of past introgression inferred with ABC (Figure 5, Table S7) are reconciled along the phylogenetic species tree in this manner, a minimum of four introgression events between ancestral lineages are retained, each of them placed in between consecutive speciation events (Figure 7, grey arrows). Accordingly, gene flow across species boundaries may have been most pronounced during early phases of divergence and since then may have decreased substantially.

Thus, the history of speciation between the contemporary high-divergence lineages may be interpreted as two-phased processes, consisting of an ancient phase with relatively strong introgression, followed by a recent and ongoing phase of much weaker or zero effective introgression. This is also suggested by the ABC model choice results, because they strongly supported introgression models over isolation models, but when we subsequently competed models that differed in the timing of introgression, past introgression was generally more probable than recent and ongoing introgression (Figure 5.) The absent or weak D-statistics (introgression younger than at most 2,000 generations, see above) between most of these lineages are also consistent with this interpretation. Hence, it appears that reproductive barriers did not arise instantly but gradually, being initially lower and then increasing substantially over time, with the genomes now largely protected against introgression. How and why initially weak reproductive barriers increased in strength remains to be investigated. One hypothesis is that divergence at many loci of small RI effect accumulated slowly (allowing for susbtantial introgression), through drift was well as adapation, until reaching a threshold beyond which RI increased abruptly to nearly perfect levels (strongly reducing introgression), a process called ‘genomic congealing’ (Feder et al. 2014; Flaxman et al. 2014). This hypothesis involves epistatic interactions between many loci of small effect, and one would expect that a severe loss of fitness occurs in a very large proportion of recombinant hybrids (e.g. F2 or back-crosses), as apparent in *Populus alba - P. tremula* hybrid zones (Christe et al. 2016). If, however, few loci of large effect control the bulk of RI, a larger fraction of recombinant hybrids should recover high fitness, which would argue against ‘genomic congealing’ as an explanation for gradual increase in RI over time. These predictions could be tested by comparing the fitness of different *Nepenthes* hybrid generations.

Sequence divergence and probabilities for ongoing gene flow allow to compare *Nepenthes* against the speciation continuum in animals identified by (Roux et al. 2016). Consistent with animal species pairs of similar divergence levels, the five high-divergence *Nepenthes* species pairs range in the ‘grey zone of speciation’, where cases of both ongoing and absent introgression are reported (Roux et al. 2016). The low-divergence *Nepenthes* in the *N. rafflesiana* complex are placed well outside this grey zone, among animal species pairs that exclusively contained signatures of ongoing gene flow. Intriguingly, the transition from low and moderate to high levels of RI may thus coincide with similar net genome-wide sequence divergence in both plants and animals.

Our genomics analyses reveal that *Nepenthes* species boundaries vary from near-complete to “semi-permeable” (Harrison and Larson 2014). The low-divergence species in the latter category, i.e. the sympatric Bornean N. *rafflesiana* complex, showed evidence for ongoing introgression, which raises questions about the factors contributing to the maintenance of species in the face of gene flow, or in other words, about the mechanisms underlying RI (Baack et al. 2015). Although the inflorescences of all three incipient species are morphologically very similar, some degree of pre-zygotic isolation likely exists, though interspecific mating is currently not quantified. Dobzhansky-Muller incompatibilities are a classical concept of postzygotic intrinsic RI mechanisms, but recent work suggests that they would be quickly lost unless they cause complete RI (Lindtke and Buerkle 2015), and none of the species studied here have achieved complete RI, as we have shown. Instead, we hypothesize that post-zygotic, extrinsic factors, i.e. ecology, could play a prominent role in *Nepenthes* RI, as has been invoked for many other plants (Abbott 2017).

As an experimental test of this hypothesis, we predict that *Nepenthes* hybrids may have low survival and little reproductive success. If ecology is indeed mediating postzygotic RI in *Nepenthes*, its strength is predicted to depend on environmental conditions (Schluter 2000; Nosil 2012). Consistent with this hypothesis, natural hybrids tend to be more common in disturbed habitats, such as logged forests, roadsides and landslides, than in intact vegetation (pers. obs.; Table S5; Peng & Clarke 2015). Reciprocal transplant experiments of parental species and hybrids in different habitats – primary vegetation versus disturbed sites – would provide an experimental test of this hypothesis (Favre et al. 2017).

Among the ecological traits that could be targets of divergent selection and drive post-zygotic RI in *Nepenthes*, the carnivorous syndrome is a prime candidate, because *Nepenthes* depend on prey capture in traps to sustain growth and reproduction and the efficiency of these traps depends on species-specific adaptations (Moran and Clarke 2010). Traps of F1 generation hybrids, however, have intermediate characteristics (Clarke 1997) and may be associated with reduced fitness. This hypothesis is supported by the pronounced pitcher trait divergence between the sister species *N. hemsleyana* and *N. rafflesiana* t.f. (Figure 6) which is much higher than expected under neutrality. Earlier work identified some of these morphological, ecological and physiological trait differences as adaptive (Clarke et al. 2011; Grafe et al. 2011; Schöner et al. 2013; Lim et al. 2015; M.G. Schöner et al. 2015; C.R. Schöner et al. 2015). In particular, the upper part of the pitchers is highly diverged between these two species. In *N. hemsleyana*, which is coprophagous, this structure serves to acoustically attract mutualistic bats that rest in the traps during the day and defecate into the traps. In contrast, *N. rafflesiana* t.f. is strictly insectivorous and traps have an associated morphology (M.G. Schöner et al. 2015). Similar ecological species differences related to divergent nutrient acquisition strategies (Moran and Clarke 2010; Thorogood et al. 2018) and leaf economy (Osunkoya et al. 2008) might contribute to RI between the remaining sympatric *Nepenthes* species.

An interesting question that our study can unfortunately not conclusively resolve, is whether the introgressing species of the *N. rafflesiana* complex in Borneo are approximating drift-migration-selection equilibrium (Lenormand 2002), or, alternatively, are far from equilibrium because of very recent speciation or secondary contact (initiation of gene flow). If these species are at equilibrium, a strong case could be made for divergent selection maintaining species boundaries in the face of gene flow. However, in case they are out of equilibrium a role for divergent selection would not be evident, and the species could be on a trajectory to speciation reversal. However, we argue that speciation times in the *N. rafflesiana* complex in Borneo are not very recent but on the order of thousands of generations, because of their wide and overlapping distribution ranges. Furthermore, our results suggest that the onset of gene flow is not recent relative to the speciation time, because both models with ongoing gene flow, ‘continuous migration since speciation’ and ‘secondary contact’, were consistent with all three species pairs. For fish species with initially similar genetic divergence as within the *N. rafflesiana* complex, speciation reversal occurred within only a few dozen generations (Vonlanthen et al. 2012). Thus, we cautiously propose that the three sympatric species in the *N. rafflesiana* complex had ample time to collapse into a hybrid swarm, but have failed to do so.

This case study uncovered that in sympatric *Nepenthes* communities, both recently and much more strongly diverged species co-occur side by side. The maintenance of species with shallow divergence despite ongoing gene flow is evidenced by pronounced pitcher trap differences that are of ecological relevance. Gene flow has occurred further in the past between lineages that have now reached more advanced stages of speciation.

Future work must determine how common sympatric communities with more than two simultaneously introgressing species are in nature, and whether systematic causes underlie the discrete occupation of the speciation continuum in local communities.

## Materials and Methods

### Sampling design and field sites

Natural populations of sympatric *Nepenthes* were sampled in 2013 at three locations in Borneo (Brunei, 4°36’N 114°35’E; Binyo Penyilam Conservation Area; Sarawak, Malaysia, 2°54’N 113°24’E; Kuching, Sarawak, Malaysia, 1°20’N 110°19’E) and in Singapore (c. 1°21’N 103°48’E). Where possible, 10 to 20 individuals from each of the three to seven locally most abundant *Nepenthes* species were sampled (Figure 1): *N. ampullaria* (Borneo and Singapore), *N. bicalcarata* (Borneo), *N. mirabilis* (Borneo), *N. gracilis* (Borneo and Singapore), *N. hemsleyana* (Borneo), and *N. rafflesiana* (Singapore). For *N. rafflesiana* in Borneo we distinguished between the “typical form” (*N. rafflesiana* t.f.) and the “giant form” (*N. rafflesiana* g.f.), as informally described by Clarke (1992, 1997). Morphospecies determination follows current taxonomy (Cheek and Jebb 2001; Scharmann and Grafe 2013).

Most samples were collected in primary peat swamp or heath forest, except for Singapore, where only secondary vegetation was available. All species were sampled from mixed stands along the same transects. Individuals of each species were sampled at a minimum distance of 50 m to avoid potential small-scale genetic structure and clonality. Plants were well established, large and climbing or with extensive rosettes. Leaf material was individually stored in nucleic acid preserving buffer solution (Camacho-Sanchez et al. 2013).

### ddRAD-seq

DNA was extracted using silica-column kits (Nucleospin Plant II, Macherey Nagel, Düren, Germany). The woody leaves were ground to fine powder with a mortar and pestle cooled by liquid nitrogen. ddRAD libraries were produced as described by (Peterson et al. 2012) with slight modifications. In brief, genomic DNA was restriction-digested with two enzymes (EcoRI and TaqI), and a 5-base barcode annealed to the EcoRI overhang. Libraries were then multiplexed to pools of 84 samples, and four pools sequenced for 101 bp single-end reads on the Illumina HiSeq 2000/2500 High-Output v4. Genotyping followed a dDocent pipeline (Puritz et al. 2014) modified for single-end reads, and specific genotype-missingness filters were applied in each analysis. In addition to the Southeast Asian *Nepenthes*, we included in the *de novo* assembly available ddRAD data of *N. pervillei*, a distantly related species from the Seychelles (H. Luqman & Ch. Küffer). Inclusion of this outgroup improved overall data quality. Further details are provided in Supporting Information S1.

### Genetic population structure

We tested for a correspondence between morphospecies and genetic groups on a per-location basis. Nucleotide sites were filtered for minimum read depth three and only polymorphic sites were retained. Principal Component Analyses were conducted with smartPCA (Price et al. 2006), and plotted in R (R Core Team 2014), based on sites present in all samples per location. Neighbor-joining trees were built from pairwise Kimura distances using rapidNJ (Simonsen and Pedersen 2011) for sites present in 90% of samples per location.

### Inference of hybrid status, parentage and class

To identify hybrids, we first ran fastSTRUCTURE (Raj et al. 2014) on a simple prior with k=2 and k=3 for each pair of morphospecies in each of the four locations (site filtering: minimum read depth three, present in >80% of samples per morphospecies, invariant sites removed, one single random SNP per RAD-tag). Samples which were not perfectly assigned to a single cluster, i.e. with >1% probability for a second cluster, were classified as putative hybrids. For some hybrids the parentage was ambiguous because of imperfect assignment in several different pairwise fastSTRUCTURE comparisons. This is probably due to shared ancestry between the potential parent species rather than hybridization between multiple species. We set the second species with the greatest minor percentage of assignment as the true parent. Results for fastSTRUCTURE k=3 helped to narrow down the list of potential parents, because the algorithm could identify the admixture component as a third cluster rather than only the two species in consideration.

We then used NewHybrids (Anderson and Thompson 2002; Wringe et al. 2017) to assign hybrids to classes: F1, F2, or the two first generation back-crosses in both directions. Samples of both parent species were assigned *a priori* as ‘pure’ (as shown by the fastSTRUCTURE results). Because NewHybrids is computationally limited to a few hundred markers, we made 100 runs each using 500 randomly sub-sampled SNPs, and then averaged the assignment probabilities from these runs. MCMC was allowed a burn-in of 1000 steps and data collected for another 1000 steps. The large number of markers and the prior identification of pure samples led independent MCMC runs to converge even after few steps. All hybrids were excluded from further analyses.

### Population phylogeny and descriptive statistics

Genotypes at minimum read depth three were filtered for presence in at least 70% of individuals in each of the 22 populations, yielding a set of 1,788 universally shared RAD-tags, which constituted a total of 177,679 sites (12,366 SNPs and 165,313 invariant sites). A concatenated supermatrix was used to estimate a maximum-likelihood (ML) tree in RAxML 8.2.4 (Stamatakis 2014) with the GTRCAT substitution model. The root of this tree was oriented as in a larger phylogeny of *Nepenthes* (Scharmann et al, in preparation). Custom scripts were used to calculate nucleotide diversity within each population (Nei and Li 1979), and for all pairs of populations the fixation index, Fst, (Weir and Cockerham 1984) and absolute sequence divergence, dxy (Nei and Li 1979).

### Tests for differential introgression in species pairs occuring sympatrically at two or more locations

We tested for introgression in sympatry using Patterson’s D-statistic (Durand et al. 2011), configured for an unrooted phylogenetic tree of two species with two geographically defined sub-populations. If no introgression occurred, the species are equally genetically different at both locations (equal count of ABBA and BABA site patterns), and the D-statistic is zero. If introgression occurred at one of the locations, or at both locations but with different magnitudes, the species may share more alleles at one location than the other (such that an inbalance of ABBA and BABA site patterns arises), and the D-statistic can deviate from zero. For each test, we selected from our dataset the species and sub-populations such that there were two locations at which two species were sympatric. Within our sampling scheme of eight species and four locations (22 populations in total), 43 suitable test combinations were possible, each conforming to the population phylogeny ((species 1 - location 1, species 1 - location 2),(species 2 - location 1, species 2 - location 2)). For each of these 43 combinations, we filtered the data for sites with minimum coverage three and for genotype presence in at least 80% of samples per population. Patterson’s D was calculated by a custom script derived from pyRAD (Eaton 2014), and the standard deviation of D was approximated by 10k bootstraps from the available sites to calculate a Z-score and two-sided p-values.

### Coalescent modelling and ABC

The general design of our simulation and ABC pipeline closely followed the approach pioneered by Roux *et al*. (2013). Principal free parameters of all models were the population sizes (Ne), the split time (speciation time), times of start and stop of migration, and migration rates. Population sizes of the ancestral and derived populations were all independent, as were the rates of bi-directional migration with their timing (i.e. asymmetric). Although we were primarily interested in detecting gene flow, we included in our models three additional processes, because they might confound the detection of migration if unaccounted for. First, changes in Ne over time (demographic events) were modelled by assuming two phases: an ancient phase of constant Ne following the split of the ancestor, followed by a phase of exponential growth or decay (demographic non-equilibrium). This added six additional free parameters (for each daughter population: ancient Ne, time of transition to recent phase, growth rate during recent phase). Given our priors, Ne could vary up to 200-fold during the time since speciation. Second, we modelled variation in Ne along the genome (Ne-heterogeneity), which is ubiquitous in natural populations due to linked or background selection (Charlesworth 2009). Following the implementation of (Roux et al. 2016), between 0 and 100% of loci were assumed to have a specific Ne below the nominal Ne. The locus-specific Ne was sampled from a beta-distribution scaled by the nominal Ne, and the two shape parameters of the beta-distribution sampled from hyperpriors. Thus, Ne-heterogeneity was modelled through five free parameters (fraction of loci with reduced Ne in each of the ancestral and both daughter populations, two hyper-parameters giving the shape of the common beta-distribution). Ne-heterogeneity was conserved through temporal changes in Ne. Third, we modelled heterogeneity in migration rates along the genome, a conceptual implication of introgression and an important diagnostic property (Roux et al. 2013; Roux, Fraisse, et al. 2014). It reflects varying strength of selection against migrant alleles among loci, i.e. a porous barrier to gene flow. Each locus was assigned specific migration rates for the two directions of migration, which were sampled from beta-distributions, scaled by the nominal migration rates. Heterogeneity in migration thus comprised four free hyper-parameters (two shape parameters for each of the two independent beta-distributions). We used 114 summary statistics, including several that are novel to ABC, such as the joint site frequency spectrum or allele frequency spectrum (Gutenkunst et al. 2009).

The sampling scheme of the real data (numbers of samples and RAD-tags, and sequence length) was closely reproduced in simulations, i.e. the same numbers of samples and RAD-tags, and the same sequence lengths. For each of the 54 sympatric species pairs, we retained 1,324 - 12,861 RAD-tags (filters: minimum read depth six, presence in at least 75 or 80% of individuals per population, excluding hybrids, excluding RAD-tags with within-population heterozygote excess, retaining monomorphic RAD-tags). After obtaining 20,000 simulated datasets under each of the four models and large, flat (uninformative) priors for all parameters, the observed data were classified by ABC model choice using the Random Forest algorithm (RF-ABC, R package abcrf, Pudlo *et al*. 2016). We devised a three-step hierarchical procedure of pairwise competitions, starting with SI against an aggregate, ‘general introgression’ model (combined PSC, OSC and CM models, weighted 2:1:1).

Second, we competed PSC against an aggregate model ‘ongoing introgression’ (combined OSC and CM models, weighted equally), and third we competed OSC against CM. Further details and a flowchart are presented in Supporting Information S2. The posterior probabilities of model choice were plotted against net sequence divergence, the average number of substitutions per site between species minus within-species polymorphisms d = dxy - (piA + piB)/2 (Nei and Li 1979).

### Pst–Fst comparison

Phenotypic data for *N. hemsleyana* and *N. rafflesiana* t.f. from Brunei were taken from Lim *et al*. (2015) who sampled the same populations as this study. We tested whether phenotypic divergence (Pst) between these sister species deviated from expectations for quantitative traits under neutrality (Leinonen et al. 2013). Pst is analogous to Qst but is estimated without a breeding experiment. We estimated Pst = (c/h^2^ * variance_between_species) / (c/h^2^ * variance_between_species + 2 * variance_within_species), where the variance components are the residuals from a linear regression model (R: lm), h^2^ is the overall proportion of variance from additive genetic effects (heritability), and c is the proportion of variance from additive genetic effects across species (Brommer 2011). As the phenotypes came from a natural common garden (Brunei), we assumed c=h^2^=1, and addressed the robustness of this assumption by calculating the critical c/h^2^ ratio (Brommer 2011). The calculation of Fst followed Weir & Cockerham (1984), and was based on 17 *N. hemsleyana* Brunei and 18 *N. rafflesiana* t.f. Brunei, i.e. the same individuals as in the rest of this study. Hybrids were excluded and individuals and genotypes filtered as in the ABC approach but monomorphic RAD-tags were removed and negative Fst estimates set to zero.

The difference between Pst and Fst was calculated using a custom bootstrap procedure in R, by bootstrap resampling from the observed Pst distribution (n = 94 phenotypic measurements, i.e. 94 pitchers were measured) and the observed Fst distribution (n = 10,397 RAD-tags) at equal sample sizes for both (n = 94), and then calculation of the difference in means; 10k replicates. Pst was considered significantly greater than Fst if the bootstrapped Pst–Fst distribution excluded zero, and the critical c/h^2^ ratio was below 0.2 (Brommer 2011).

## Acknowledgements

We thank Abdul Hadzid Harith Tinggal, Ieney Daud and their family for hospitality in Brunei. In Sarawak, Ch’ien C. Lee, Pearl Ee, and Joanes Unggang are thanked for logistics and help with field work. Lam Weng Ngai is thanked for assistance with field work in Singapore. Hirzi Luqman and Christoph Küffer are thanked for sampling in the Seychelles. Permissions were kindly granted by the Ministry of Industry and Primary Resources of Brunei Darussalam, the Agri-Food and Veterinary Authority and the National Parks Board of Singapore, the Forest Department of Sarawak (Malaysia), and the Seychelles Agricultural Agency. We thank Claudia Michel for wet lab support. The Genetic Diversity Center Zürich, the Quantitative Genomics Facility Basel, and the Functional Genomics Center Zürich provided sequencing and computational support. Keygene N.V. owns patents and patent applications protecting its Sequence Based Genotyping technologies. J. Stapley, K. Csilléry, and two anonymous reviewers provided helpful comments on earlier versions of the manuscript. This work was supported by the ETH Zürich and the Rübel foundation.

## Data Accessibility

- exact sampling coordinates are withheld for conservation reasons
- demultiplexed Illumina sequencing reads will be uploaded to NCBI SRA
- genotypes in .vcf format and bioinformatic scripts produced for this study will be uploaded to DRYAD respectively github.com/mscharmann

## Author Contributions

MS carried out field work, bioinformatics and statistical analyses, and some of the molecular lab work; FM and TUG contributed logistic support and materials; MS and AW conceived and designed the study, and wrote the manuscript.

